# EnrichViz: An Interactive R Shiny Application for Visualization of Pathway Enrichment Results from Omics Data

**DOI:** 10.64898/2026.06.19.733398

**Authors:** Rolando Garcia-Milian

## Abstract

Pathway and functional enrichment analysis is a cornerstone of omics data interpretation, enabling researchers to map differentially expressed proteins or genes onto curated biological processes, signaling cascades, and molecular functions. While tools such as Ingenuity Pathway Analysis (IPA), g:Profiler, and Enrichr are widely used to generate ranked enrichment results, translating these tabular outputs into clear, publication-ready figures remains a time-consuming step that typically requires custom scripting and familiarity with visualization libraries — a significant barrier for researchers without a computational background.

Here we present EnrichViz, a self-contained, browser-based R Shiny application that enables interactive, code-free visualization of pathway and functional enrichment results from quantitative proteomics, transcriptomics, and metabolomics experiments. EnrichViz accepts three standard CSV files as input — a normalized abundance matrix, a sample annotation or metadata file, and enrichment results from any platform that exports tabular output — and produces six complementary, publication-ready visualizations: bar and bubble plots for ranking enriched terms by significance, chord diagrams for exploring pathway-molecule connectivity, clustered heatmaps for displaying Z-score normalized expression patterns across experimental groups, and boxplots or violin plots for examining the abundance distribution of individual proteins, genes, or metabolites. The application supports both raw p-values and pre-transformed -log10(p) values through automatic detection, and all plot parameters are adjustable in real time through a graphical sidebar. Every figure can be exported as a high-resolution PNG file at 300 dpi. EnrichViz is implemented in R using the Shiny, ggplot2, pheatmap, and circlize packages, and is freely available at https://rgmilian.shinyapps.io/EnrichViz/.

## Introduction

High-throughput omics experiments — including quantitative proteomics and bulk transcriptomics — routinely produce lists of hundreds to thousands of differentially expressed proteins or genes. Interpreting these lists in a biologically meaningful way requires pathway and functional enrichment analysis, which maps individual molecules onto curated biological processes, signaling cascades, and molecular functions. Tools such as Ingenuity Pathway Analysis (IPA, Qiagen), g:Profiler (Kolberg et al., 2023), and Enrichr (Kuleshov et al., 2016) among others, are widely used for this purpose and produce ranked tables of enriched terms alongside the molecule lists that drive each enrichment.

However, translating tabular enrichment results into clear, publication-ready figures remains a time-consuming step that typically requires custom scripting in R or Python, familiarity with visualization libraries, and repeated manual adjustment of plot parameters. Researchers without a computational background may find this step to be significant barrier, and even experienced users must re-implement similar code across projects.

EnrichViz was developed to address this gap. It is a self-contained, browser-based R Shiny application that accepts three standard CSV files as input — normalized abundance data, metadata or sample annotation, and enrichment results — and produces four complementary, interactive visualizations without requiring any programming. All plot parameters are adjustable in real time through a graphical sidebar, and every figure can be downloaded as a high-resolution .png file suitable for publication or presentation. EnrichViz is designed to be tool-agnostic: it accepts enrichment output from any platform that exports a .csv table, and it supports both raw p-values and pre-transformed -log10(p) values through automatic detection. The application can be downloaded from GitHub (https://github.com/rologmilian/enrichviz.git) and run locally or accessed freely at https://rgmilian.shinyapps.io/EnrichViz.

## Components

### Implementation

EnrichViz is implemented entirely in R (≥ 4.2.0) using the Shiny framework (Chang, 2026) for the graphical user interface and it is freely available online at https://rgmilian.shinyapps.io/EnrichViz.Visualization is handled by four packages: ggplot2 (part of the tidyverse) (Wickham H, 2019) for the bar plot, bubble plot, and boxplot and violin plot; pheatmap (Kolde, 2019) for clustered heatmaps; and circlize (Gu et al., 2014) for chord diagrams. Data wrangling is performed with dplyr and tidyr from the tidyverse. The complete application is contained in a single *app.R* file and requires no database connection or internet access once installed.

### Input data

The application requires three comma-separated value (CSV) files, all with a header row.

#### Normalized abundance data

A matrix in which rows correspond to proteins, genes or metabolites, and columns correspond to individual samples, with one additional column containing unique molecule identifiers (e.g. gene symbols or protein accessions). Values are expected to be normalized (e.g., normalized LFQ intensities, VST-normalized counts, TMT reporter ion ratios) since no further normalization is applied by the application.

#### Metadata or Sample annotation

A minimum of two-column table mapping each sample identifier to an experimental group or condition. Sample identifiers must exactly match the column names in the abundance or count matrix.

#### Enrichment results

A table produced by a pathway or functional enrichment tool (e.g. IPA, g:Profiler, Enrichr, or similar) in which each row represents one enriched term. The table must contain at minimum: a column with the pathway or function name, a column with a significance metric (p-value or -log10(p-value)), and a column listing the molecule identifiers that drive the enrichment for each term, separated by a user-specified delimiter (comma, slash, semicolon, or pipe). Optional numeric columns such as gene count or overlap ratio can be used as additional visual encodings in the bubble plot.

### Enrichment p-value handling

The application automatically determines whether the selected significance column contains raw p- or p-adjusted values (values in the range 0–1) or values already transformed to the -log10 scale (values > 1). In the former case, a -log10 transformation is applied before plotting; in the latter, values are used as-is. The detected transformation and the column used are reported in the status line of the *Bar / Bubble Plot* tab.

### Visualizations

#### Bar / Bubble Plot

Pathways are ranked in descending order by their -log10(p-value) and the top N terms (default 20, range 5–200) are displayed. In bar plot mode, each pathway is represented by a horizontal bar whose length encodes significance. In bubble plot mode, the X-axis retains the same significance ranking while bubble area is mapped to a second user-selected numeric column (e.g. gene count or overlap size) and bubble fill color intensity is mapped to a third column (e.g. False Discovery Rate or fold enrichment) using a continuous color scale from one of ten available palettes (Blues, Reds, Purples, Greens, OrRd, YlOrRd, RdYlBu, viridis, magma, plasma). When the color column contains raw p-values, a -log10 transformation is applied automatically, and the legend label is updated to reflect this. Bubble diameter is scaled to a user-defined range (default 3– 15 points) (Figures 1A, 2A, 3A).

**Figure 1.**
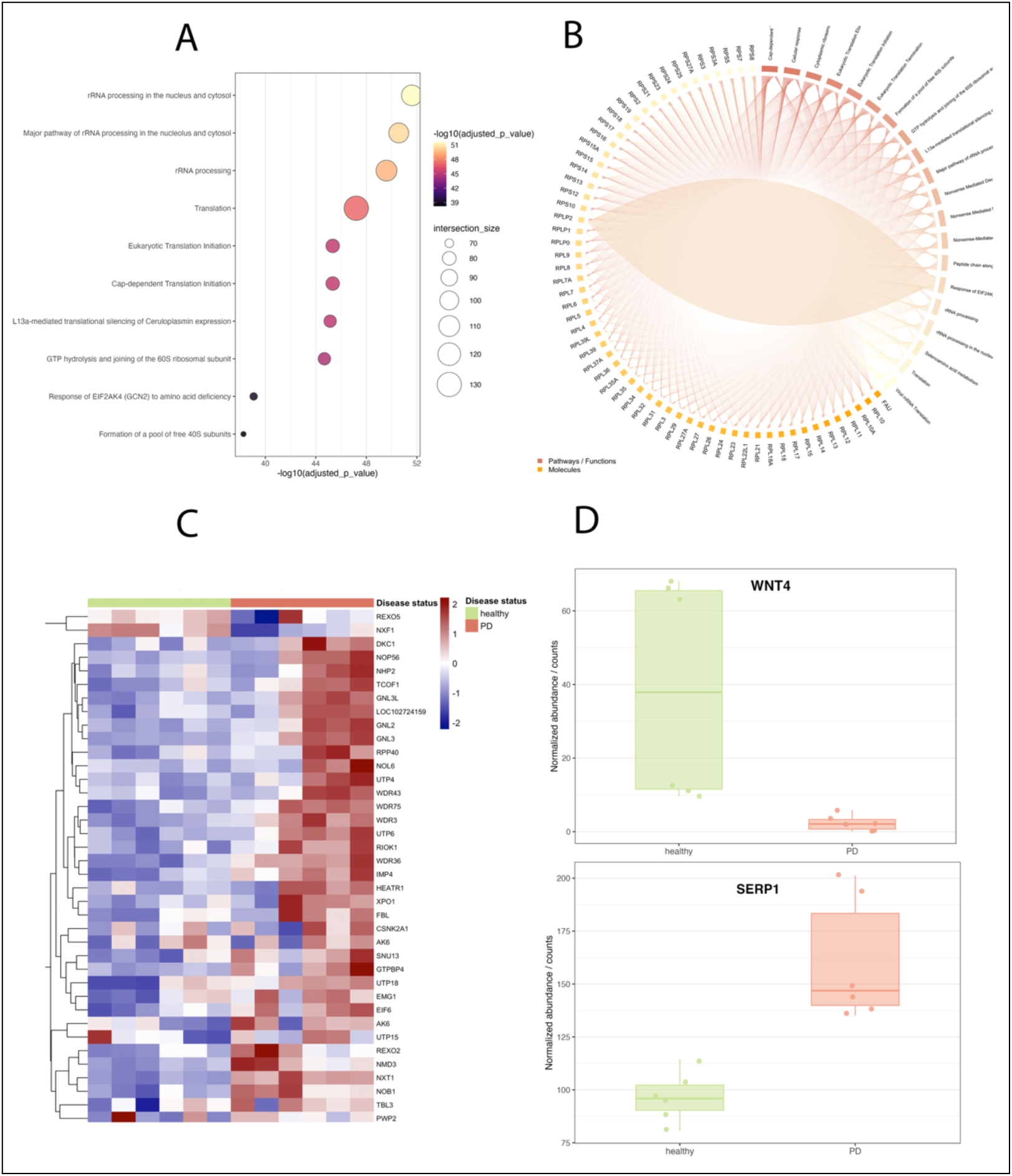
Visualization of overrepresentation analysis on g:ProfilerGOSt of bulk RNA-seq dataset GSE185009 of neuronal progenitor cells derived from discordant monozygotic twins with Parkinson’s Disease compared with healthy controls. A) Bubble plot of top 10 enriched pathways (KEGG, Reactome, WP). B) Chord plot of the top 20 pathways and proteins with 4 or more connections. C) Hierarchical clustering heatmap of proteins overlapped with pathway Ribosome biogenesis in eukaryotes. D) Boxplots of protein WNT4 (upper panel) and SERP1 (lower panel) in healthy and Parkinson’s Disease groups.

**Figure 2.**
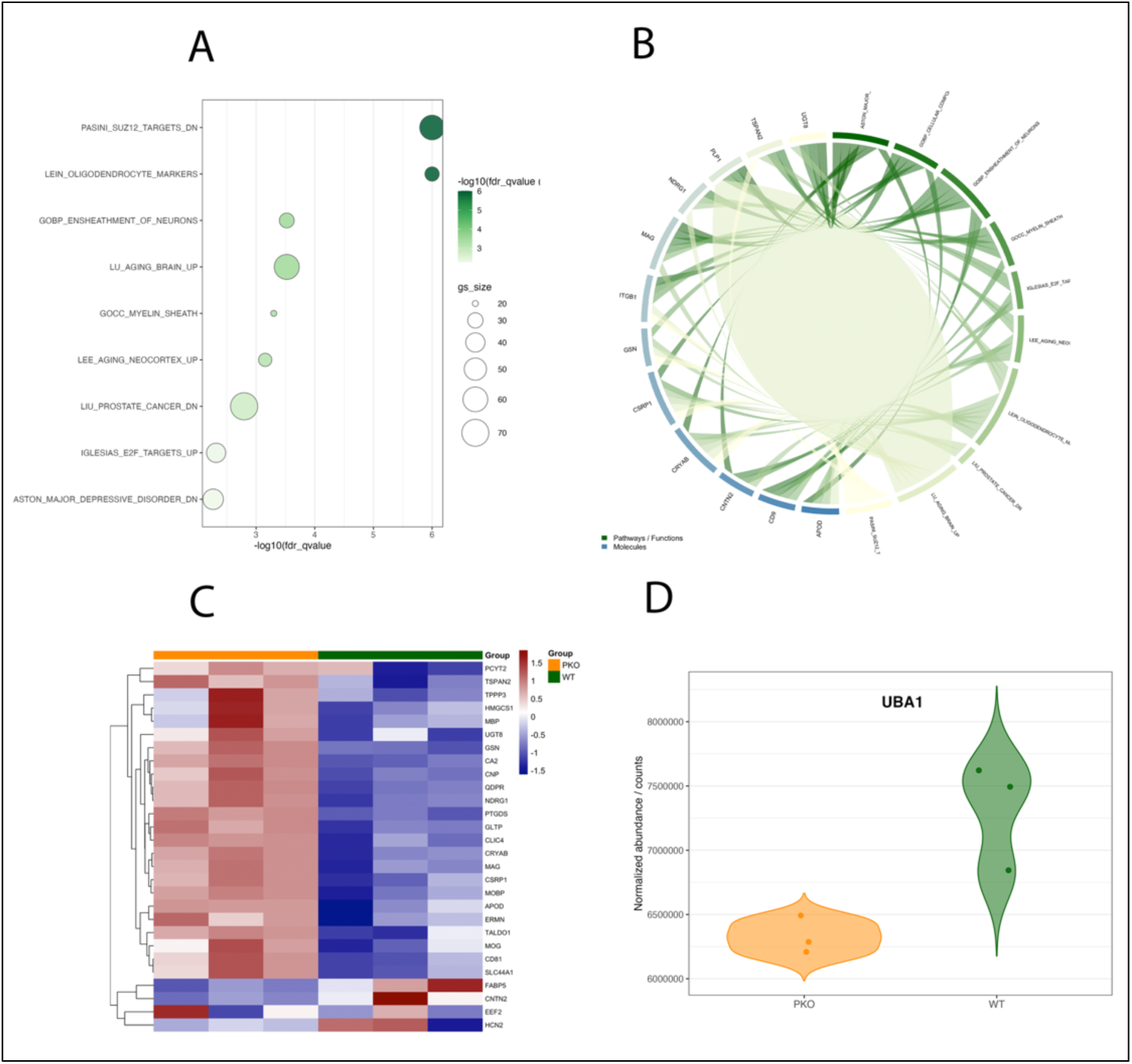
Visualization of proteomics enrichment analysis data. A) Bubble plot of top 10 enriched molecular database signatures from a Gene Set Enrichment Analysis comparing protein abundances from PKO vs WT. B) Chord plot of the top signatures and proteins with 7 or more connections. C) Hierarchical clustering heatmap of proteins overlapped with LEIN_OLIGODENDROCYTES_MARKERS signature. D) Boxplots of protein UBA1 abundance in PKO and WT groups

**Figure 3.**
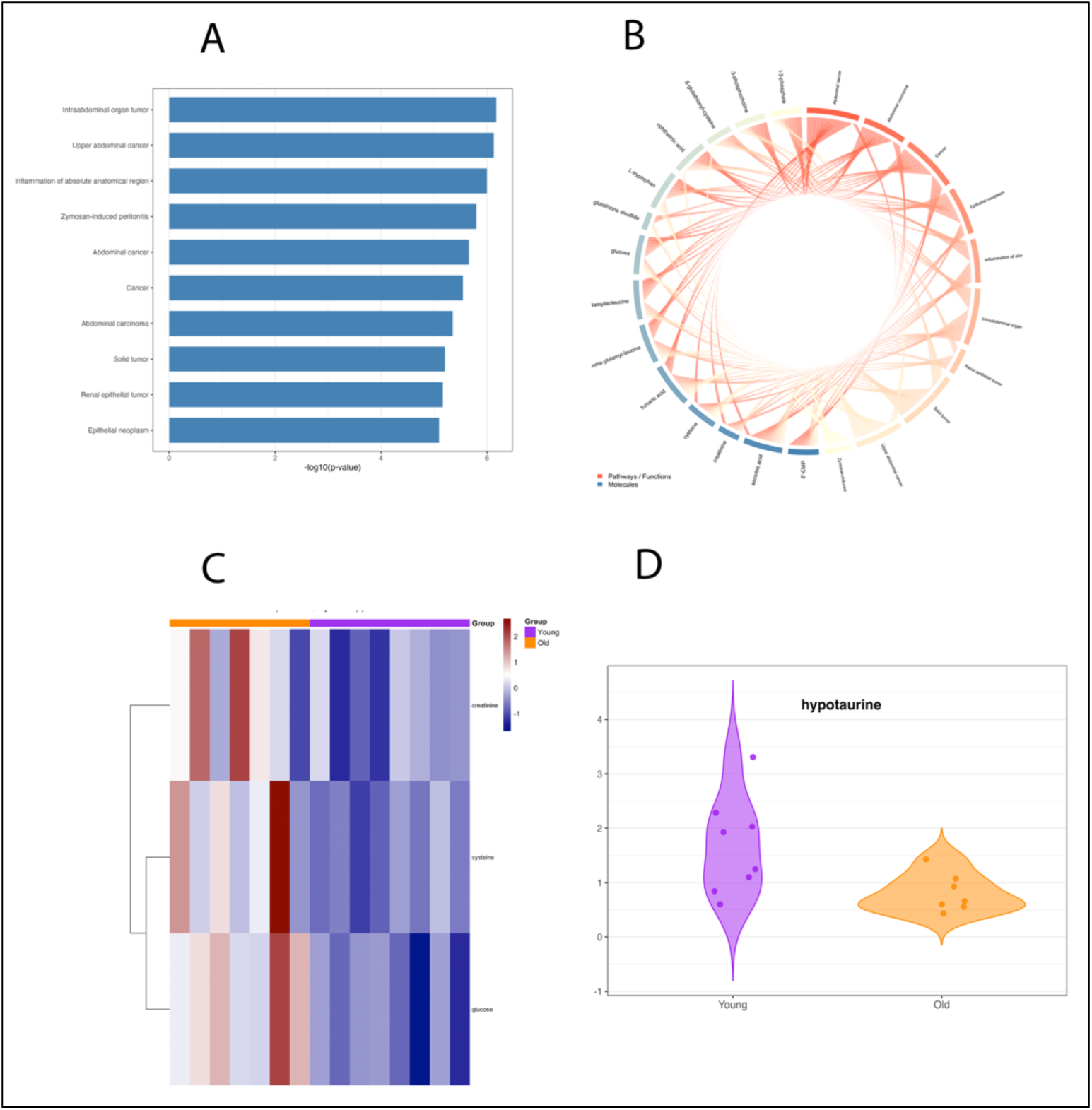
Visualization of metabolomics enrichment analysis data from a Core Analysis (Ingenuity Pathway Analysis) of differentially abundant metabolites comparing livers from 2-years-old versus young mice. A) Bar plot of top 10 enriched diseases and biofunctions. B) Chord plot of the top enriched Ingenuity biofunctions and metabolites with 2 or more connections. C) Hierarchical clustering heatmap of metabolites overlapped with Inflammation of organ. D) Violin plot of metabolite hypotaurine in young and old groups.

#### Chord Diagram

The top N pathways by significance (default 10) and the molecules they contain are represented as sectors on a circular axis. Each connection between a pathway sector and a molecule sector is drawn as a ribbon whose width is proportional to the number of shared connections. Pathway sectors are colored using a gradient derived from a user-specified base color (default tomato red) and molecule sectors from a separate gradient (default steelblue); color shade within each family reflects alphabetical order only and encodes no biological variable. Molecules that appear in fewer than a user-defined number of pathways (default 2) are excluded to reduce visual clutter. Label font sizes and the inner radius of the diagram are adjustable (Figures 1B, 2B, 3B). The chord diagram is rendered using the *chordDiagram()* function from the circlize package. (Gu et al., 2014)

#### Heatmap

For a user-selected pathway or function, the normalized abundance values of all matched proteins are extracted, Z-score scaled row-wise (Mean = 0, Standard Deviation = 1 per protein) and displayed as a clustered heatmap. Rows (proteins) are hierarchically clustered using the default complete-linkage method with Euclidean distance as implemented in pheatmap; columns (samples) are ordered by experimental group but not clustered. A color annotation bar above the heatmap indicates group membership for each sample using user-specified colors. The color scale runs from dark blue (low) through white (zero) to dark red (high). Row labels are displayed when 100 or fewer proteins are present (Figures 1C, 2C, 3C). All heatmaps for all enriched terms can be exported as a ZIP archive in a single click.

#### Boxplot / Violin Plot

For a single user-selected protein or gene, the raw normalized abundance values are displayed grouped by experimental condition. In boxplot mode, each group is represented by a box showing the median, interquartile range, and whiskers extending to 1.5 × IQR (Interquartile Range; individual sample values are overlaid as jittered points and outlier symbols from the box geometry are suppressed to avoid double-plotting. In violin plot mode, the box is replaced by a mirrored kernel density estimate (*geom_violin(), trim = FALSE*) with individual points overlaid. Group colors are shared with the heatmap annotation and are configurable through the sidebar (Figures 1D, 2D, 3D).

### Output

All four visualizations can be downloaded as PNG files at 300 dots per inch (dpi). Bar and bubble plots are exported at 10 × 8 inches; chord diagrams at 10 × 10 inches; individual heatmaps at 8 × 10 inches; boxplots and violin plots at 8 × 6 inches. Download buttons are available both in the sidebar and below each plot in the main panel. The exported filename reflects the active plot type (e.g., *Barplot_Enriched_Pathways.png* vs. *Bubbleplot_Enriched_Pathways.png*; *Boxplot_<protein>.png* vs. *Violin_<protein>.png*).

## Availability

EnrichViz is written in R and released under the MIT License. The source code is available at the project repository GitHub (https://github.com/rologmilian/enrichviz.git). A hosted instance is available at https://rgmilian.shinyapps.io/EnrichViz/. Local deployment requires R ≥ 4.2.0 and the following CRAN packages: shiny, tidyverse, pheatmap, and circlize.

## Datasets

Considering that enrichment analysis methods (e.g., Overrepresentation Analysis (Tavazoie et al., 1999), Gene Set Enrichment Analysis (Subramanian et al., 2005)) rank results based on statistical significance, it is strongly recommended to filter the results of the enrichment or pathway analysis for biological relevance before generating visualizations. This requires ample knowledge of the biological context and the phenotype being studied. This section contains the three different datasets (e.g., transcriptomics, proteomics, metabolomics) used to test the EnrichViz app. All datasets used in this manuscript including normalized, metadata, and enrichment results can be found at https://doi.org/10.60600/YU/F4MKC3 (Garcia-Milian, 2026)

### Visualization of bulk RNA-sequencing enrichment results

Normalized counts and metadata from a Parkinson’s Disease (PD) study (Vlasov et al., 2021) was downloaded from GREIN database https://www.ilincs.org/apps/grein/?gse=GSE185009. This dataset contains bulk RNA expression profiling of Induced Pluripotent Stem Cells (IPSC) and Neuronal Progenitor Cells (NPC) Derived from discordant monozygotic twins with Parkinson’s Disease. DESeq2 differential analysis (Love et al., 2014) comparing NPC PD samples versus NPC healthy samples was done on the GEO Datasets GEO2R platform https://www.ncbi.nlm.nih.gov/geo/geo2r/?acc=GSE185009. A total of 3805 differentially expressed genes (DEG) were obtained with adjusted p < 0.05. DEG were used for Overrepresentation Analysis on g:Profiler GOSt (https://biit.cs.ut.ee/gprofiler/gost, version e114_eg62_p19_27110d83) (Kolberg et al., 2023) with following parameters: sources KEGG, REAC, WP; organism, hsapiens; significance threshold method, g_SCS. Functional Annotation chart cutoOs were EASE 0.1, and count threshold 5 (Figure 1).

### Visualization of proteomics Gene Set Enrichment Analysis (GSEA) results

For visualization of proteomics data, we used published LC-MS/MS label-free proteomics data from a study on the use of methamphetamine addictive behaviors in young adult male rats. We compared Parkin knockout PKO (*Park2*^*−/−*^) versus wild type samples (Sharma et al., 2025). Normalized protein abundances were loaded into the Gene Set Enrichment Analysis (version 4.4.0) desktop application and GSEA (Subramanian et al., 2005) was run with the following parameter: random seed 1649521726881, nperm 1000, set_max 500, set_min 15, metric Signal2Noise, permute gene_set. A subset of enrichment results was used for visualization purposes (Figure 2).

### Visualization of metabolomics enrichment results

The results of a metabolomics study comparing livers from young and 2-year-old mice was downloaded (Houtkooper et al., 2011). Changes in normalized metabolites intensities between old and young livers were assessed for each metabolite by either Student’s t-test or Welch’s t-test on RStudio (version 4.5.2). A total of 46 differentially expressed metabolites (p-value < 0.05) was functionally analyzed on Ingenuity Pathway Analysis with the Core Analysis tool. Enriched biofunctions were downloaded and used for visualization (Figure 3).

Detailed instructions for the installation and deployment of EnrichViz are available in the EnrichViz GitHub repository https://github.com/rologmilian/enrichviz.git.

## Acknowledgements

This work was supported in part by the Yale/NIDA Neuroproteomics Center through the National Institutes of Health grant P30 DA018343.

